# Deep-Palm: An integrated deep learning framework for structure-aware prediction of protein S-palmitoylation

**DOI:** 10.64898/2026.03.05.709753

**Authors:** Maolin Deng, Jiaming Huang, Han Wang, Shenghong Fu, Wei Wang, Liyi Li, Yu-Jian Kang, Bo Xu

**Affiliations:** Chongqing Key Laboratory of Intelligent Oncology for Breast Cancer, Intelligent Oncology Innovation Center Designated by the Ministry of Education, School of Medicine, Chongqing University, Chongqing 400030, China; Chongqing Key Laboratory of Intelligent Oncology for Breast Cancer, Intelligent Oncology Innovation Center Designated by the Ministry of Education, Chongqing University Cancer Hospital and Chongqing University School of Medicine, Chongqing 400030, China

## Abstract

Protein S-palmitoylation is a reversible lipid modification that regulates protein localization, trafficking, and signaling. Its dysregulation has been implicated in cancer and therapeutic resistance, making accurate site annotation important for understanding disease-related regulatory mechanisms. However, experimental identification of S-palmitoylation sites remains labor-intensive, highlighting the need for computational tools that can support large-scale candidate-site prioritization. S-palmitoylation-site recognition requires the integration of multiple levels of information surrounding candidate cysteine residues, including sequence context, local protein properties and structural features. Here, we present Deep-Palm, a deep learning framework that integrates four complementary branches: amino acid sequence, physicochemical properties, ESM embedding and spatial structure. On the testing set, Deep-Palm achieved an AUC of 0.950 and outperformed the compared predictors, including pCysMod, MusiteDeep and GPS-Palm. Deep-Palm also showed stable performance across diverse cysteine sequence contexts and Gene Ontology functional groups. Feature-level analyses revealed distinct spatial structural and protein property patterns between palmitoylated and non-palmitoylated peptide windows. Independent mass spectrometry datasets further demonstrated the ability of Deep-Palm to sensitively identify previously unannotated S-palmitoylation sites. Together, Deep-Palm provides an accurate and biologically informative framework for S-palmitoylation-site prediction, facilitating the discovery of novel candidate S-palmitoylation sites.

## Introduction

The functional diversity of the eukaryotic proteome is substantially expanded by post-translational modifications (PTMs)^[1]^. Among diverse PTMs, S-palmitoylation is a reversible lipid modification that acts as a dynamic regulatory mechanism for protein activity, localization, and interaction networks^[2]^. S-palmitoylation involves the formation of a labile thioester bond between a 16-carbon saturated fatty acid, palmitate and the thiol group of specific cysteine residues^[3]^. This process is regulated by the opposing activities of two enzyme families: zinc finger DHHC-domain-containing protein acyltransferases (ZDHHCs), which catalyze palmitate addition, and acyl-protein thioesterases (APTs), which mediate depalmitoylation^[2]^. Since S-palmitoylation is reversible, it can rapidly modulate membrane residence and signaling output of proteins in response to upstream cues, and its dysregulation has been implicated in a variety of diseases, particularly cancer^[4-6]^.

Several public resources curate experimentally supported S-palmitoylation site annotations, such as SwissPalm and CysModDB^[7-9]^. Although the number of annotated S-palmitoylation sites has steadily increased, current annotations remain incomplete due to the experimental complexity and limited throughput of enrichment- and chemical-reporter-based methods^[2,10,11]^. Therefore, accurate computational prediction of S-palmitoylation sites is essential for systematic prioritization of candidate cysteine residues and large-scale discovery of previously unannotated sites^[12-14]^.

Several computational tools have been developed to predict palmitoylated cysteines using machine learning or deep learning methods, such as GPS-Palm^[12]^, MusiteDeep^[15]^ and pCysMod^[14]^. These methods typically represent each cysteine by a fixed-length sequence window and predict palmitoylation with sequence composition features. Another typical strategy utilizes sequence similarity with known palmitoylated peptides. For example, CSS-Palm 2.0 applies clustering-and-scoring strategy in which query peptides are classified according to similarity to known palmitoylated clusters^[13]^. PalmPred uses PSI-BLAST-derived PSSM profiles for the evaluate the palmitoylation probability^[16]^. And MDD-Palm scans palmitoylation substrate motifs for prediction^[17]^.

However, sequence composition features such as k-mer frequency are often difficult to interpret biologically, as they do not explicitly capture higher-order secondary or tertiary structure and physicochemical properties associated with S-palmitoylation recognition. Meanwhile, methods based on similarity to previously known sites may be inherently biased by existing database annotations, thereby limiting the discovery of novel S-palmitoylation events, especially in non-model organisms. Moreover, recent protein language models have demonstrated powerful capability in encoding protein evolutionary, structural and functional properties directly from amino acid sequences. Although these representations provide rich contextual information for PTM and significantly increases recognition accuracy, they have not yet been integrated into S-palmitoylation site-level prediction models^[18,19]^.

Thus, we present Deep-Palm, a multi-view deep learning framework for S-palmitoylation site prediction. Deep-Palm integrates amino acid sequence, spatial structure, protein physicochemical properties and ESM protein language model embeddings. In evaluation with multiple eukaryotic species, Deep-Palm accurately predicts S-palmitoylation sites across diverse sequence contexts and Gene Ontology functional categories. Feature-level analyses reveal distinct structural and physicochemical patterns associated with S-palmitoylation, suggesting that Deep-Palm captures biologically relevant determinants underlying palmitoylation-site recognition. Independent mass spectrometry (MS) data demonstrate that Deep-Palm effectively identifies previously unannotated sites. Collectively, Deep-Palm provides a framework for systematic annotation and discovery of S-palmitoylation sites, enabling further investigation into the underlying mechanisms of protein palmitoylation.

## Materials and Methods

### Data preprocessing strategy

To construct a high-quality dataset for S-palmitoylation-site prediction, we collected experimentally supported S-palmitoylation annotations from SwissPalm^[7]^, CysModDB^[8]^ and the GPS-Palm^[12]^ training dataset, which was curated from experimentally verified S-palmitoylation records reported in previous studies. Positive samples were defined as 31-residue peptide windows centered on experimentally verified S-palmitoylated cysteine residues. Negative samples were defined as cysteine-centered windows from cysteine sites with no reported S-palmitoylation annotation in any of the curated data sources used in this study.

To reduce homology bias and potential information leakage, redundancy among positive sequences was removed using CD-HIT at a 60%sequence identity threshold^[20]^. After positive-sample redundancy removal, the positive set contained 3,445 samples. An equal number of negative samples was then randomly selected to construct a balanced dataset containing 6,890 samples, including 3,445 positive samples and 3,445 negative samples. The balanced dataset was then divided into training and independent test sets using stratified sampling, with 10%of the data reserved for independent testing while preserving the class distribution across splits. This split yielded 6,201 training samples and 689 independent test samples.

### Feature calculation

#### 1. Evolutionary Semantic Encoding

To encode contextual sequence representations, we used precomputed ESM-2 (3B) embeddings for each cysteine-centered peptide window^[21]^. Each 31-residue window was represented as an ESM-2 embedding matrix. The embedding matrix was transposed to a channel-first format before convolutional processing. The ESM embedding branch processed the channel-first embedding matrix using two vConv-based one-dimensional convolutional blocks, followed by adaptive max pooling and a fully connected classifier^[22]^. In this branch, vConv applies a learnable mask to the convolutional kernel, allowing the effective receptive field to be adjusted during training^[22]^.

#### 2. Structure-aware graph representation

To incorporate structure-derived information, predicted structures of the 31-residue peptide windows were generated using ESMFold^[23]^.

For each peptide window, residues were represented as graph nodes, and spatial contacts were used to define graph edges. A spatial contact was defined when the residue-residue distance based on Cα-Cα coordinates was less than 8 Å^[24]^. Each peptide window was encoded as a 6×31×31 residue-pair tensor, including residue-residue distance based on Cα-Cα coordinates, minimum heavy-atom distance, side-chain-center distance, binary spatial contact, long-range contact, and normalized sequence separation^[25-28]^.

For residues i and j in a peptide window of length L, the residue-pair feature vector was defined as follows:

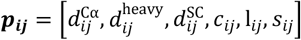

where *p*_*ij*_ denotes the residue-pair feature vector for residues i and j, and L denotes the length of the peptide window. 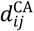 denotes the residue-residue distance calculated from the Euclidean distance between the Cα atom coordinates of residues i and j. 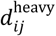 denotes the minimum Euclidean distance between any non-hydrogen atom in residue i and any non-hydrogen atom in residue j. 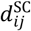 denotes the Euclidean distance between the side-chain centers of the two residues. The side-chain center was defined by the Cβ atom when available; otherwise, it was calculated as the mean coordinate of side-chain non-hydrogen atoms, with the Cα coordinate used as a fallback. *c*_*ij*_ denotes the binary spatial-contact indicator, which was set to 1 when 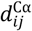 was less than 8 Å and to 0 otherwise. *l*_*ij*_ denotes the long-range spatial-contact indicator, which was set to 1 when residues i and j were separated by at least six positions in the sequence and also satisfied the spatial-contact criterion. *s*_*ij*_ denotes the normalized sequence separation, calculated as the absolute sequence-position distance between residues i and j divided by *L* − 1.

Stacking this feature vector over all residue pairs yielded a 6×31×31 residue-pair tensor for each peptide window. The residue-pair tensor was first processed by a 2D convolutional module, and node-level representations were obtained by aggregating the resulting pair features along the two residue dimensions. A two-layer GCN was then used to propagate information over the contact-derived adjacency matrix^[29]^. An identity matrix was added to the contact adjacency matrix before normalization. The adjacency matrix was then normalized by symmetric degree normalization, and the resulting matrix was averaged with its transpose to ensure symmetry before graph convolution.

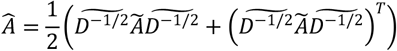

where Ã = A + I denotes the adjacency matrix after adding the identity matrix, 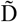 denotes the degree matrix of Ã, and Â denotes the final normalized adjacency matrix obtained after symmetric degree normalization and transpose averaging. Graph convolution was then applied as:

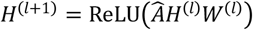

where H(l) is the node representation at layer l and W(l) is the learnable weight matrix of the corresponding GCN layer.

The resulting graph representation enabled the structure branch to aggregate information from residues that may be distant in the primary sequence but spatially close in the predicted structure.

#### 3. Protein properties modeling

The protein properties branch represented each residue in the 31-residue peptide window using 14 AAindex-derived protein-property descriptors^[30]^. These descriptors included hydropathy index, polarity, volume, flexibility parameter, alpha-helix frequency, beta-sheet frequency, isoelectric point, hydrophilicity value, residue accessible surface area, B-value-derived flexibility, membrane propensity, beta-turn frequency, average reduced side-chain distance, and relative partition energy. For each 31-residue peptide window, these descriptors formed a 31×14 residue-level protein property matrix:

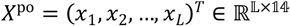

where x_t_ denotes the 14-dimensional AAindex descriptor vector at residue position t.

The descriptor matrix was first projected to a hidden representation and then processed by a bidirectional LSTM encoder^[31]^. This encoder modeled dependencies among residues across the peptide window in both sequence directions. An attention-based readout was applied to the hidden states to estimate residue-level contributions to the branch-level prediction^[32]^.

The attended representation was passed to a fully connected classifier to produce the prediction score of the protein properties branch.

#### 4. Amino acid sequence branch based on k-mer features

The amino acid sequence branch encoded each 31-residue peptide window using 2-mer, 3-mer, and 4-mer tokens. For a peptide sequence s = (a_1_, a_2_, …, a_L_), the k-mer token sequence was defined as:

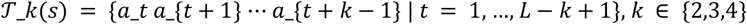

where k denotes the k-mer length and L = 31 in this study.

Separate vocabularies were built for each k value using the training sequences, and unseen k-mer tokens were mapped to an out-of-vocabulary token. For each k value, token indices were transformed into 32-dimensional embeddings and processed by a one-dimensional convolutional encoder^[33]^.

### Ensemble learning and model training

To integrate the four base branches, Deep-Palm generated two ensemble outputs: weighted blending and stacked ensemble^[34,35]^. For weighted blending, branch-level probability outputs were combined using non-negative weights selected from out-of-fold predictions on the training set. The weighted blending output was defined as:

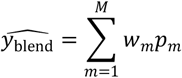

where *p*_*m*_ denotes the probability output of branch *m, w*_*m*_ denotes the corresponding blending weight, and *M* = 4 in the full Deep-Palm model. For the stacked ensemble, branch-level probabilities were used as input features for a logistic regression meta-learner trained on out-of-fold predictions^[34]^. The stacked ensemble output was defined as:

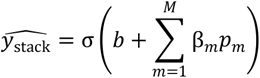

where β_*m*_ denotes the coefficient learned by the meta-learner, b denotes the intercept term, and σ (⋅) denotes the sigmoid function.

Training Protocol: The model was implemented in PyTorch^[36]^. We adopted a stratified 5-fold cross-validation scheme to generate out-of-fold predictions for ensemble training. Each branch model was optimized using the AdamW optimizer^[37]^. Weight decay was set to 5×10^-4^ for the ESM embedding, protein properties, and spatial structure branches and to 0.001 for the amino acid sequence branch. Binary cross-entropy with logits was used as the loss function. Equivalently, after sigmoid transformation of the logits, the loss was written as:

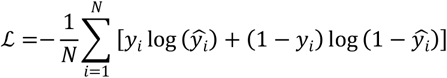

Early stopping was triggered if the Area Under the Receiver Operating Characteristic Curve (AUROC) on the validation set did not improve for 5 consecutive epochs.

### Feature-level statistical analysis

To compare structure-derived and protein-property features between palmitoylated and non-palmitoylated peptide windows, feature-level analyses were performed on the independent test set. For structure-derived features, the six ESMFold-derived residue-pair descriptors were summarized across valid residue pairs within each 31-residue peptide window. Contacts were defined using a residue-residue distance based on Cα-Cα coordinates threshold of 8 Å, and long-range contacts were defined as contacting residue pairs with sequence separation |i - j|≥6. Contact ratio at each sequence separation was calculated as the fraction of valid residue pairs that satisfied the contact criterion within the corresponding separation bin. For protein-property features, the 14 AAindex-derived descriptors were calculated at each position of the 31-residue peptide window. Sample-level group differences were assessed using the two-sided Mann-Whitney U test. For position-pair or position-wise analyses, false-discovery-rate correction was applied across tested cells, and the Mann-Whitney U statistic was converted to a signed Z-score to indicate the direction and magnitude of group differences.

### Positive and putative negative site identification in MS data

Palmitoylation MS data (biotin-enriched) were obtained from two publicized datasets, including a mouse hepatocellular carcinoma dataset (5 replicates)^[38]^ and a human HeLa cell dataset (3 replicates)^[39]^. For the mouse hepatocellular carcinoma dataset, label-free DDA MS was used for palmitoylation-site quantification with Q-Exactive HF-X mass spectrometer (Thermo Scientific). For the human HeLa dataset, DIA MS was used for palmitoylation-site quantification with Orbitrap Astral mass spectrometer (Thermo Scientific). The database searching was performed using Proteome Discoverer 2.4 against the UniProt Mus musculus or Homo sapiens reference proteome respectively. Trypsin was used as the digestion enzyme, with a maximum of two missed cleavages. The precursor tolerance was set to 10 ppm, and the MS/MS tolerance was set to 0.02 ppm. Oxidation (M), protein N-terminal acetylation, deamidation (NQ), N-ethylmaleimide modification (C) and carbamidomethylation (C) were set as variable modifications. The target-reverse strategy was used for database searching, and the false discovery rate was controlled at 0.01 at both the peptide-spectrum match and protein levels. Protein quantification was performed using razor and unique peptides.

To retrieve high-confidence palmitoylated and non-palmitoylated cysteine residues from MS data, we applied the following stringent criteria. Positive residues (palmitoylated sites) were defined as cysteine residues identified in peptides detected across all replicates with high confidence, q-value < 0.01 and Carbamidomethyl modification. To define high-confidence non-palmitoylated cysteine sites while minimizing false negatives caused by lack of protein expression, we selected cysteine residues showing no evidence of palmitoylation (palmitoylome abundance = 0) but consistently detected in the corresponding proteome datasets (proteome abundance > 0) across all replicates. The 15-residue flanking sequences (±15 amino acids) were extracted for the positive and negative cysteines (xpos and xneg). Sequences shorter than 15 residues from the terminus were padded with ‘X’ to maintain a uniform input length.

### Prediction with GPS-Palm, pCysMod and MusiteDeep

We selected three palmitoylation prediction tools that are still publicly accessible for performance comparison with Deep-Palm. Predictions were performed on the same datasets with Deep-Palm. GPS-Palm was downloaded from https://gpspalm.biocuckoo.cn/download.php and receives input sequences centered on cysteine residues with a window size of 31. The parameters were set as follows: Organism = General; Threshold = All. pCysMod was accessed through its web server at http://pcysmod.omicsbio.info and receives input sequences centered on cysteine residues with a window size of 31. The parameters were set as follows: Modification type = S-palmitoylation. MusiteDeep was accessed through its web server at https://www.musite.net and receives input sequences centered on cysteine residues with a window size of 31. The parameters were set as follows: PTM type = Palmitoylation.

### Performance evaluation

Model performance was evaluated using standard binary-classification metrics, including sensitivity, specificity, accuracy, and AUC. Palmitoylated cysteine-centered peptide windows were treated as positive instances, whereas cysteine-centered peptide windows without reported S-palmitoylation annotation in the curated data sources were treated as negative instances. Let TP, TN, FP and FN denote the numbers of true positives, true negatives, false positives and false negatives, respectively. The threshold-dependent metrics were computed as:

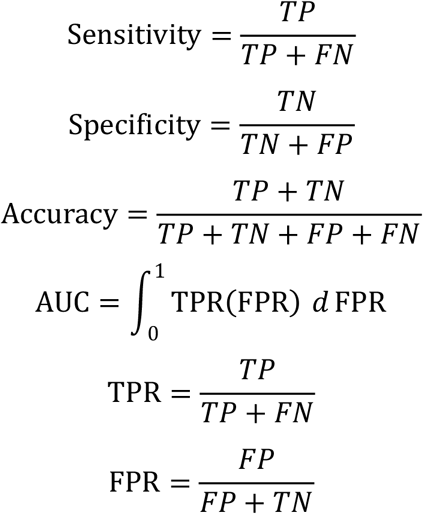

Model training and evaluation were conducted on an NVIDIA A100 GPU.

## Results

### Dataset composition and Deep-Palm architecture

To model the complex interplay among sequence, structural, and protein property determinants underlying S-palmitoylation, we developed a biologically informed multi-view framework for accurate site prediction. The curated dataset for model training and testing comprised experimentally validated S-palmitoylation sites from diverse eukaryotic species (Table 1), supporting model development across multiple species. Deep-Palm adopts a multi-branch deep learning architecture in which all feature representations are derived from a 31-amino-acid sequence window centered on the cysteine residue (± 15 aa). The framework consists of four complementary feature extraction modules: k-mer-based sequence encoding, protein property descriptors, protein language model embeddings derived from ESM-2, and spatial representations calculated from ESMFold-predicted structures (Fig. 1)^[21,30]^. Each branch independently learns latent representations associated with palmitoylation propensity using branch-specific neural encoders, including convolutional blocks for k-mer features, a BiLSTM encoder for protein property descriptors, vConv-based convolutional blocks for ESM-2 embeddings, and CNN-GCN modules for spatial structure features, followed by branch-specific prediction heads that generate site-level probabilities^[22,29,31]^. To integrate heterogeneous biological signals, branch-level outputs were combined using ensemble strategies based on weighted blending or stacked meta-learning, enabling adaptive integration of sequence-derived, structural, protein property and protein language embeddings for final prediction.

**Table 1.** Dataset Statistics.

| Organism | Total | Negative | Positive |
| --- | --- | --- | --- |
| *Mus musculus* | 3049 | 585 | 2464 |
| *Homo sapiens* | 1558 | 687 | 871 |
| *Arabidopsis thaliana* | 356 | 344 | 12 |
| *Rattus norvegicus* | 273 | 239 | 34 |
| *Caenorhabditis elegans* | 158 | 156 | 2 |
| *Saccharomyces cerevisiae* | 134 | 130 | 4 |
| *Drosophila melanogaster* | 137 | 130 | 7 |
| *Bos taurus* | 117 | 112 | 5 |
| *Xenopus laevis* | 113 | 113 | 0 |
| *Dictyostelium discoideum* | 111 | 111 | 0 |
| Others (206 species) | 884 | 838 | 46 |
| Total (all samples) | 6890 | 3445 | 3445 |

**Figure 1.**
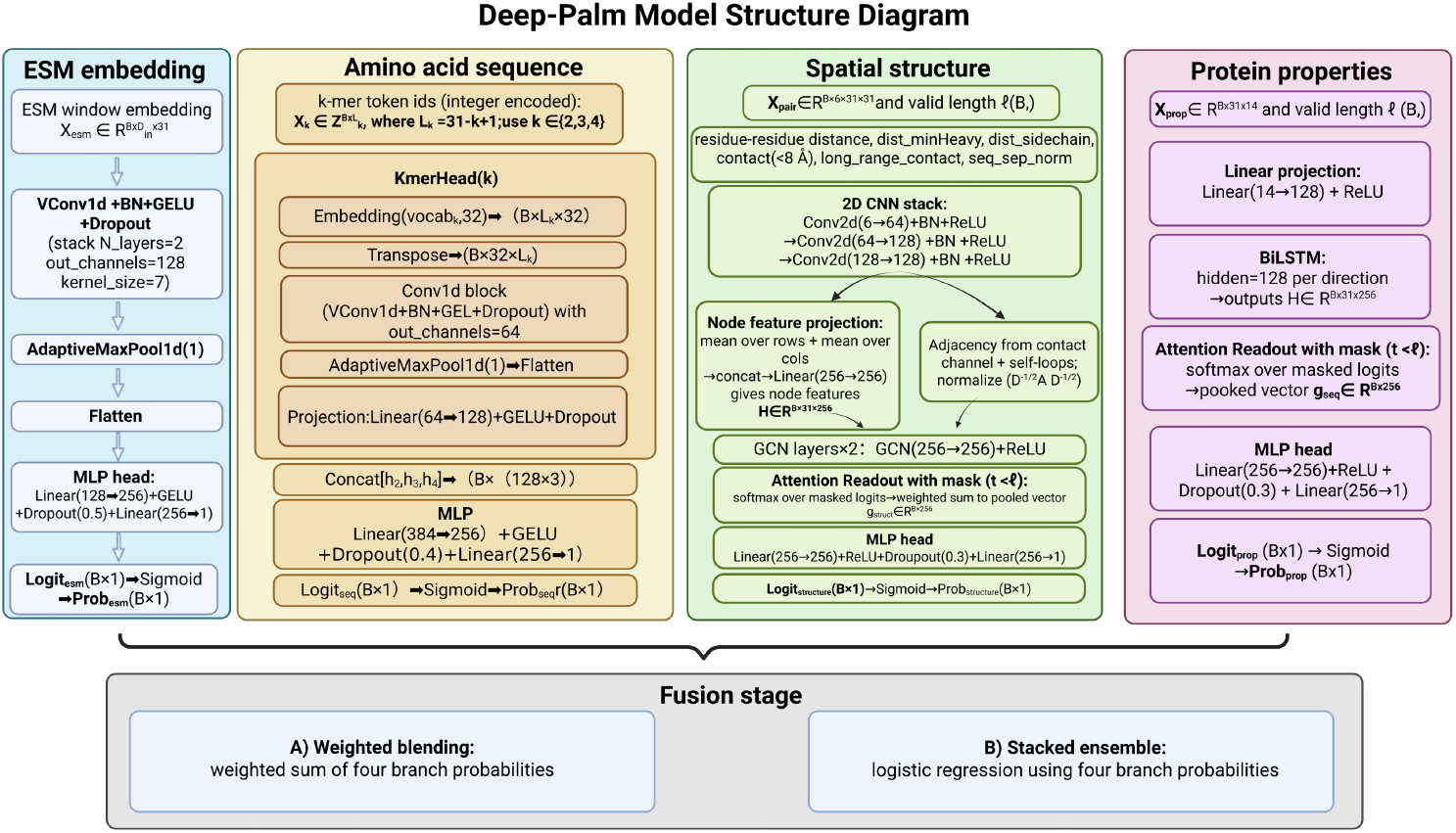
Overview of the Deep-Palm framework.

### Deep-Palm shows balanced performance in S-palmitoylation-site prediction

We first evaluated Deep-Palm on an independent test set. At the branch level, the protein properties and ESM embedding modules achieved AUC values of 0.939 and 0.934, respectively, whereas the amino acid sequence and spatial structure modules yielded AUC values of 0.882 and 0.680, respectively (Fig. 2A). Ensemble integration further improved performance, with weighted blending achieving an AUC of 0.949 and the stacked ensemble output, defined as the final Deep-Palm model, reaching the highest AUC of 0.950 (Fig. 2A, Table 2). At the selected decision threshold, Deep-Palm achieved a sensitivity of 0.85, a specificity of 0.90, and an accuracy of 0.87 (Fig. 2C).

**Table 2.**
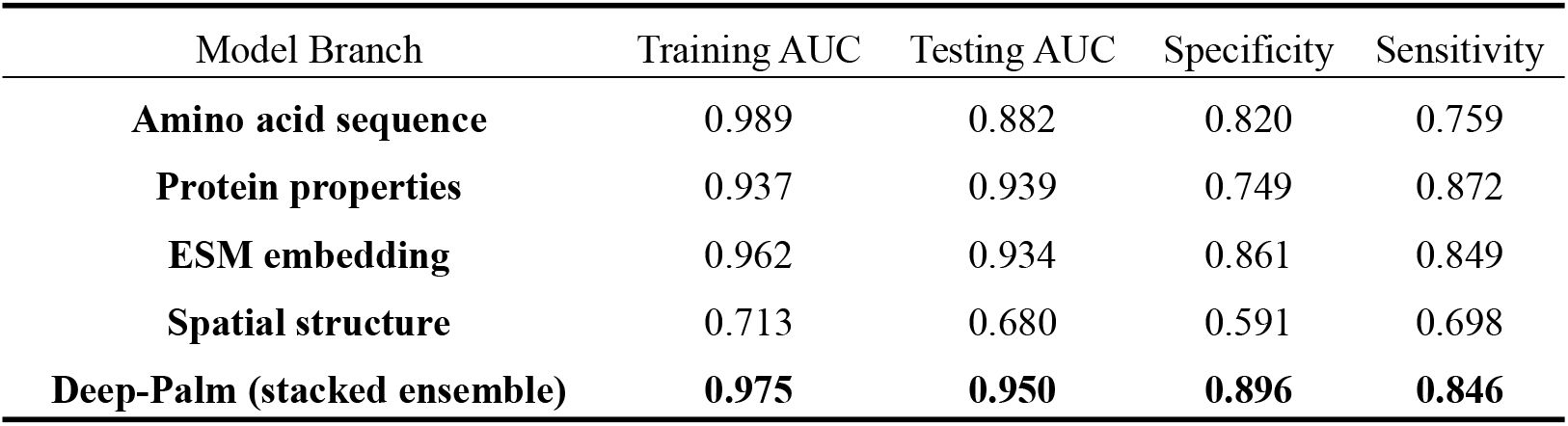
Performance comparison of individual branches and the integrated Deep-Palm model on the training set and independent test set.

**Figure 2.**
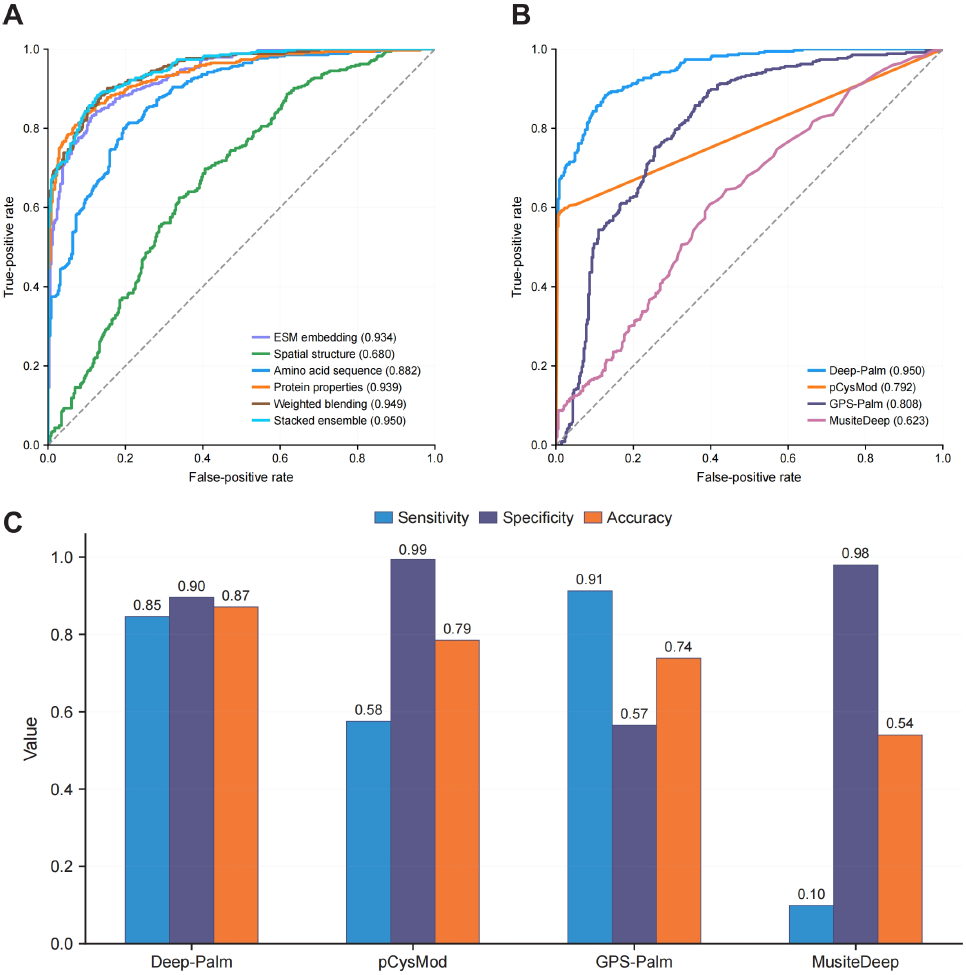
Performance evaluation of Deep-Palm and comparison with existing predictors on the independent test set. **(A)** ROC curves of the four individual branches, weighted blending, and Deep-Palm (stacked ensemble) on the independent test set. **(B)** ROC curves of Deep-Palm, pCysMod, GPS-Palm, and MusiteDeep on the independent test set. **(C)** Comparison of sensitivity, specificity, and accuracy for Deep-Palm, pCysMod, GPS-Palm, and MusiteDeep on the independent test set.

We further compared Deep-Palm with existing S-palmitoylation site predictors which are are currently accessible on the same independent test set. In the ROC analysis, Deep-Palm achieved an AUC of 0.950, outperforming GPS-Palm (0.808), pCysMod (0.792), and MusiteDeep (0.623) (Fig. 2B). Deep-Palm also maintained a balanced trade-off between sensitivity and specificity, whereas GPS-Palm showed higher sensitivity but reduced specificity, and pCysMod and MusiteDeep showed higher specificity but lower sensitivity (Fig. 2C). Overall, these results demonstrate that Deep-Palm achieved superior predictive performance while maintaining a balanced classification profile on independent data.

### Deep-Palm exhibits robust performance across diverse cysteine contexts and functional protein groups

Since variations in cysteine positional distribution, such as N- or C-terminal proximity, and local cysteine density within the sequence window may introduce potential prediction bias, we systematically assessed whether the performance of Deep-Palm was sensitive to different local sequence contexts. Specifically, the independent test set was stratified according to (i) the number of cysteine residues within the 31-residue sequence window, (ii) the distance between the target cysteine and its nearest neighboring cysteine, and (iii) the relative position of the target cysteine within the N- to C-terminal sequence region. Across these stratifications, Deep-Palm generally achieved stable predictive performance and was comparable to or higher than existing predictors in most comparisons (Fig. 3A-D), indicating that its performance is not strongly dependent on specific local cysteine density or positional configurations.

**Figure 3.**
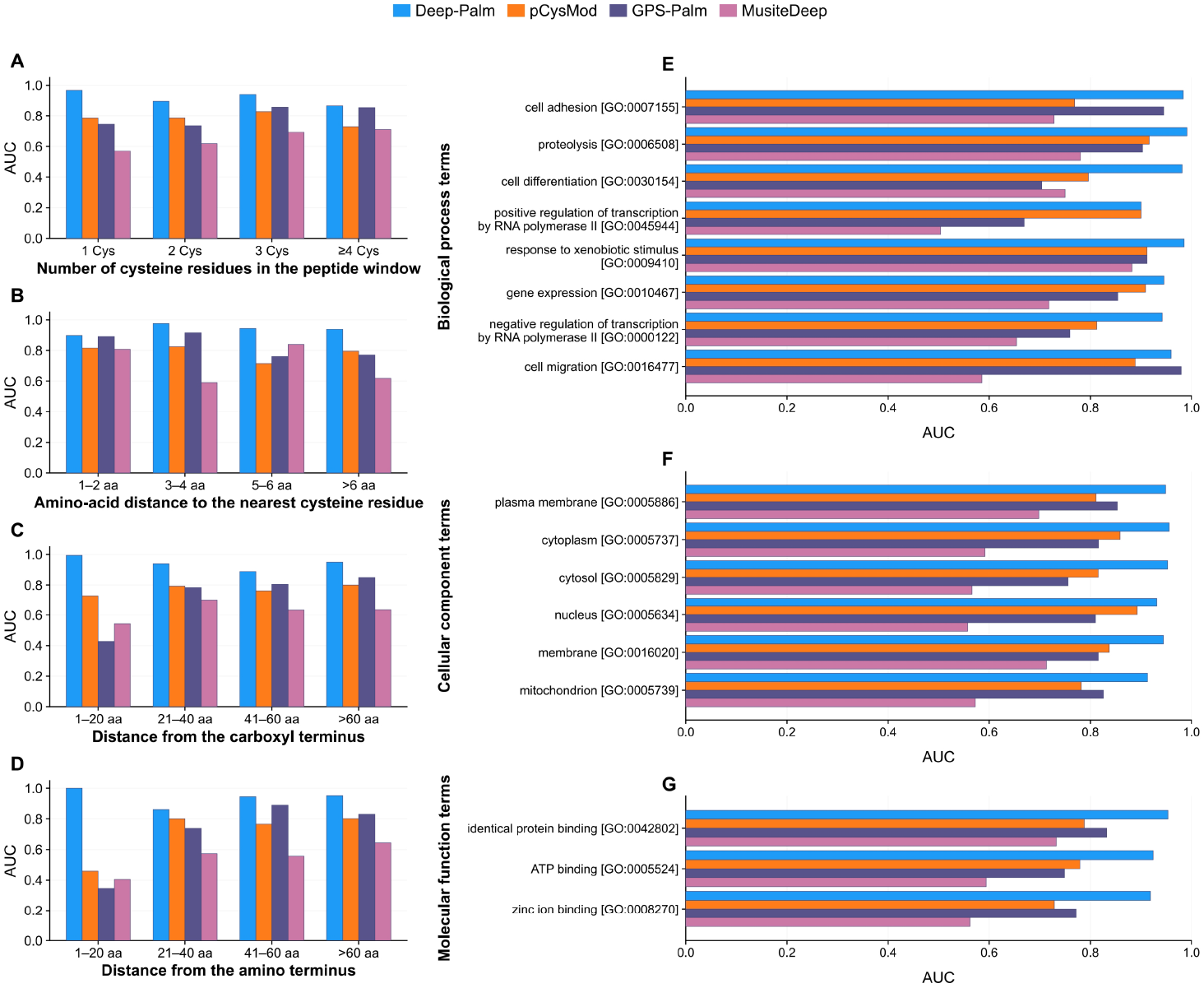
Performance of Deep-Palm and existing predictors across peptide-level cysteine contexts and Gene Ontology functional groups. **(A)** Performance across peptide groups defined by the number of cysteine residues within the 31-residue peptide window. **(B)** Performance across peptide groups defined by the distance from the target cysteine to the nearest neighboring cysteine. **(C)** Performance across peptide groups defined by the distance from the target cysteine to the peptide C terminus. **(D)** Performance across peptide groups defined by the distance from the target cysteine to the peptide N terminus. **(E)** Performance across independent-test-set subsets grouped by Biological Process (BP). **(F)** Performance across independent-test-set subsets grouped by Cellular Component (CC). **(G)** Performance across independent-test-set subsets grouped by Molecular Function (MF).

Given that S-palmitoylation is closely associated with membrane localization and protein trafficking processes, model performance may be influenced by functional and subcellular annotation biases across different protein groups. We further assessed model performance across Gene Ontology (GO) categories, including Biological Process (BP), Cellular Component (CC), and Molecular Function (MF). Deep-Palm maintained stable and high performance across GO groups, with AUC values above 0.90, and generally outperformed existing predictors (Fig. 3E-G). These results demonstrate that Deep-Palm is largely robust to functional heterogeneity and retains stable predictive performance across cysteine sites located in diverse sequence contexts.

### Spatial structure and protein property characteristics at S-palmitoylation sites revealed by Deep-Palm

Benefiting from the biologically informed feature construction in Deep-Palm, which integrates sequence, spatial structure, and protein property representations, we sought to characterize feature patterns associated with S-palmitoylation site recognition. This analysis compared palmitoylated and non-palmitoylated peptide windows and was not intended to assign causal effects to individual features. For spatial structure representations, residue-residue distance based on Cα-Cα coordinates was calculated among residues within each 31-residue peptide window using ESMFold-predicted structures. Palmitoylated peptide windows showed larger mean residue-residue distance based on Cα-Cα coordinates than non-palmitoylated peptide windows (Fig. 4A), indicating greater spatial separation among residues within the local window. Position-pair analysis further showed increased residue-residue distance based on Cα-Cα coordinates for multiple residue pairs near the two ends of the 31-residue window in palmitoylated sites (Fig. 4B), revealing that such spatially conformational patterns may contribute to S-palmitoylation site recognition.

**Figure 4.**
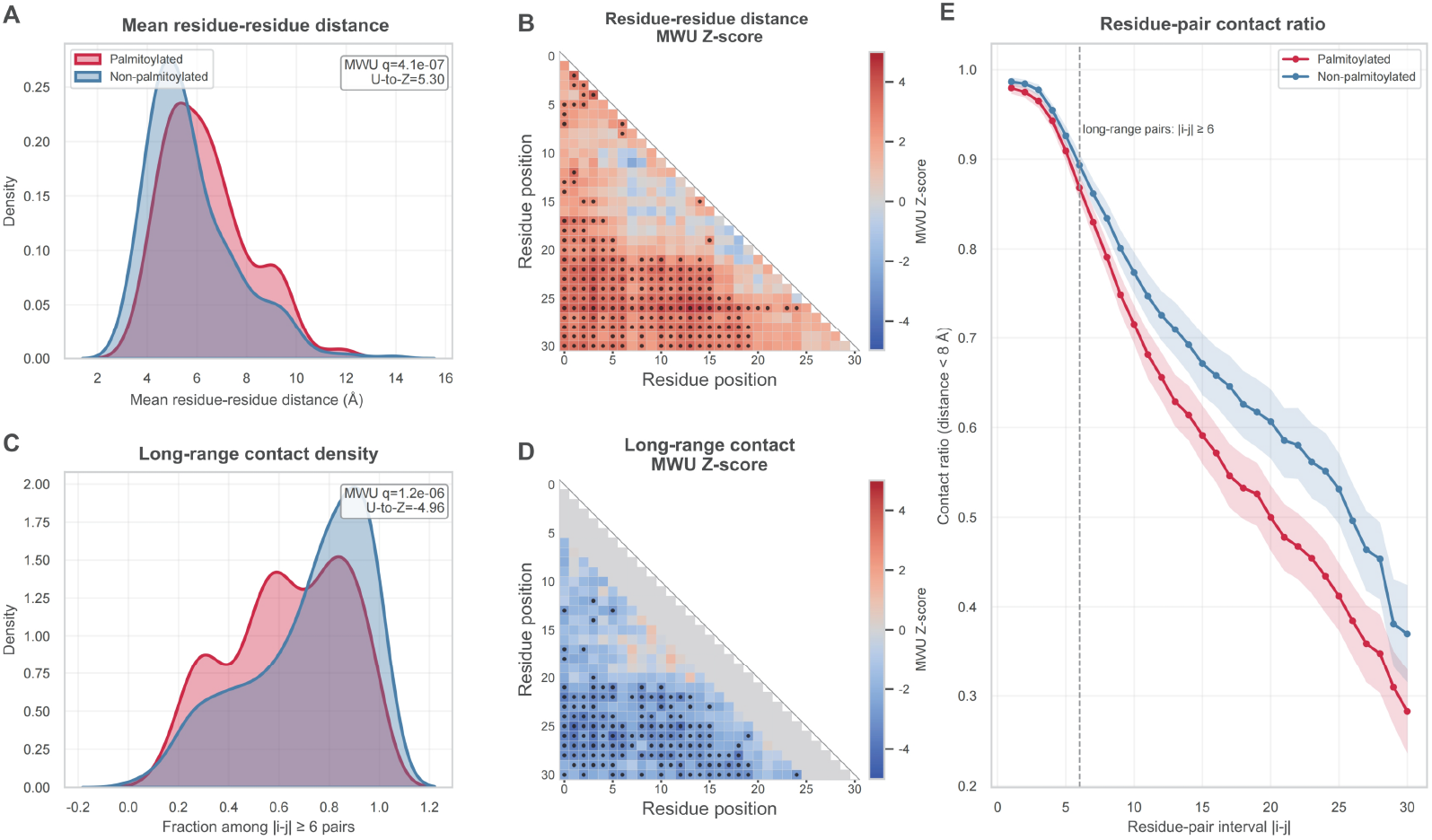
Spatial structure features at S-palmitoylation sites revealed by Deep-Palm. **(A)** Distribution of mean residue-residue distance based on Cα-Cα coordinates in palmitoylated and non-palmitoylated peptide windows. Each value represents the mean distance across valid **(B)** non-diagonal residue pairs within a 31-residue peptide window. **(C)** Lower-triangle position-pair heatmap of the signed Z-score converted from the Mann-Whitney U statistic for residue-residue distance based on Cα-Cα coordinates. Positive values indicate larger values in palmitoylated peptide windows, whereas negative values indicate larger values in non-palmitoylated peptide windows. Black dots indicate position pairs with FDR-adjusted q < 0.05. **(D)** Distribution of long-range contact density. Long-range contact density was calculated as the fraction of residue pairs with sequence separation |i - j| ≥ 6 that satisfied the spatial-contact criterion, defined as residue-residue distance based on Cα-Cα coordinates less than 8 Å. **(E)** Lower-triangle position-pair heatmap of the signed Z-score converted from the Mann-Whitney U statistic for long-range contact. Gray regions indicate residue pairs with sequence separation |i - j| <6, which were not used for long-range contact analysis. Black dots indicate position pairs with FDR-adjusted q < 0.05. **(F)** Contact ratio stratified by sequence separation. The x-axis shows the residue-pair interval |i - j|, and the y-axis shows the contact ratio, defined as the fraction of valid residue pairs satisfying the spatial-contact criterion at each interval. Lines show the mean contact ratio across peptide windows, and shaded regions indicate bootstrap 95% confidence intervals. The dashed vertical line marks the threshold used to define long-range residue pairs, |i - j|≥6.

Long-range contact density was then analyzed to assess whether residues that are distant in sequence tend to be spatially close in the predicted local structure. Long-range contacts were defined as residue pairs separated by at least six positions in the sequence and satisfying the spatial-contact criterion, defined as residue-residue distance based on Cα-Cα coordinates of less than 8 Å. Long-range contact density was calculated as the fraction of long-range contacts among eligible residue pairs within each 31-residue peptide window. Non-palmitoylated peptide windows showed higher long-range contact density than palmitoylated peptide windows (Fig. 4C), indicating more frequent spatial contacts between sequence-distant residues. Consistently, the position-pair heatmap based on the signed Z-score converted from the Mann-Whitney U statistic showed stronger long-range contact signals in non-palmitoylated peptide windows across multiple residue-pair regions (Fig. 4D). When contact ratio was stratified by sequence separation, non-palmitoylated peptide windows maintained higher contact ratios across most medium- and long-range sequence separations (Fig. 4E). Together, these results indicate that spatial structure features at S-palmitoylation sites are associated with larger mean residue-residue distance based on Cα-Cα coordinates and lower long-range contact density.

For protein properties, we calculated 14 physicochemical features using AAindex from the same 31-residue peptide windows (Fig. 5A)^[30]^. The position-wise comparison of physicochemical differences between palmitoylated and non-palmitoylated peptide windows showed distinct trends across multiple descriptors at the upstream neighboring amino acids (Fig. 5A). The upstream residues adjacent to the palmitoylated cysteines exhibited increased hydrophobicity and decreased polarity and contact energy, suggest that the physicochemical properties of the residues may play important roles in S-palmitoylation site recognition (Fig. 5A). Among the 14 protein property features, alpha-helix frequency exhibited the strongest separation between palmitoylated and non-palmitoylated peptide windows, with consistently higher values observed in palmitoylated sites (Fig. 5B), indicating that S-palmitoylation preferentially occurs in alpha-helical regions. In addition, isoelectric point (pI), hydropathy, flexibility, and hydrophilicity also showed higher values in palmitoylated peptide windows (Fig. 5C, E-G). In contrast, turn frequency showed the opposite direction, with higher values in non-palmitoylated peptide windows (Fig. 5D).

**Figure 5.**
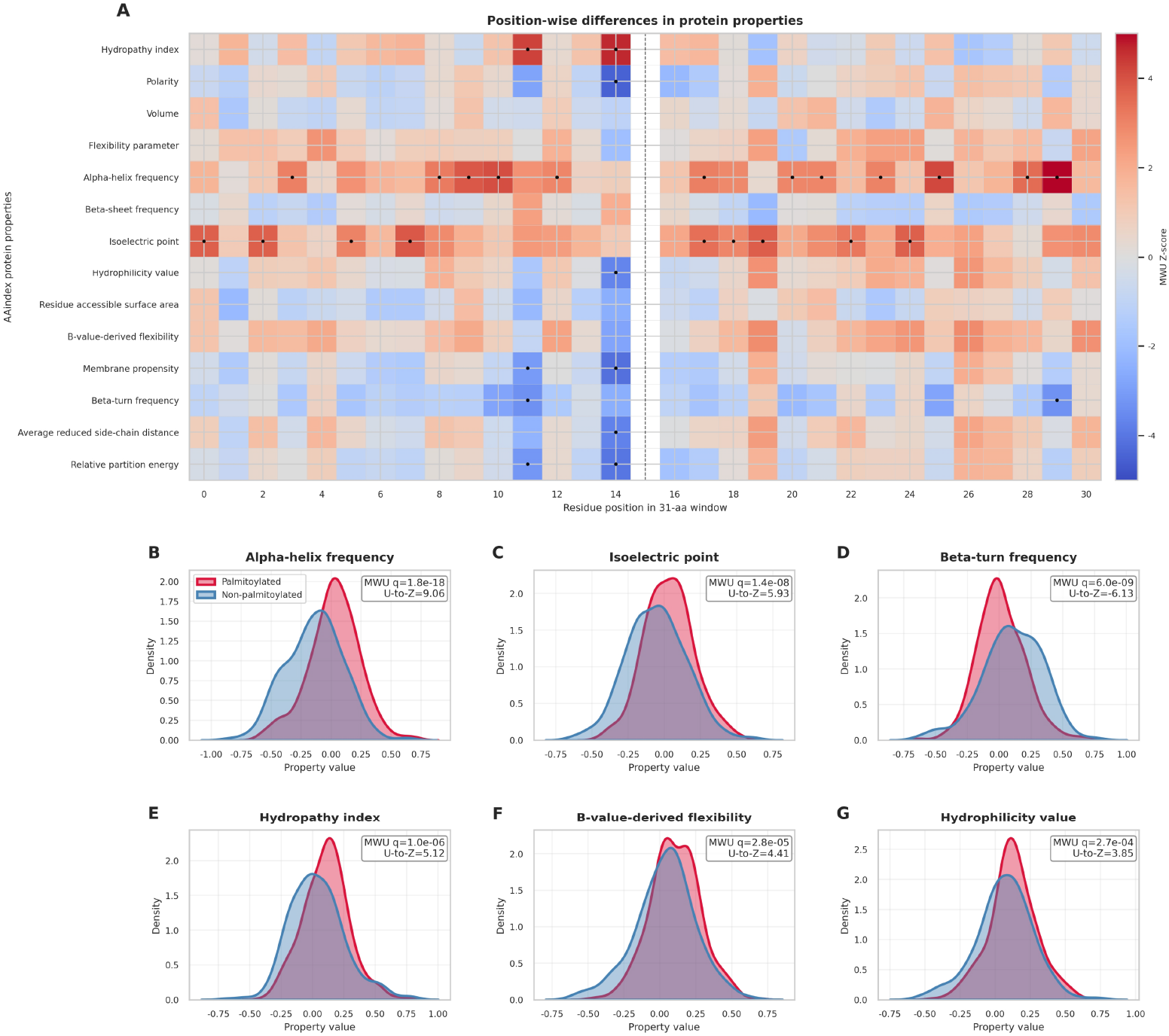
Protein property features at S-palmitoylation sites revealed by Deep-Palm. **(A)** Position-wise heatmap of the signed Z-score converted from the Mann-Whitney U statistic for 14 AAindex-derived protein property descriptors across the 31-residue peptide window. The x-axis shows residue positions in the 31-residue window, and the y-axis shows the 14 AAindex-derived protein property descriptors. Positive Z-scores indicate higher descriptor values in palmitoylated peptide windows, whereas negative Z-scores indicate higher descriptor values in non-palmitoylated peptide windows. Black dots indicate positions with FDR-adjusted q < 0.05. The dashed vertical line marks the center cysteine. **(B)** Density distribution of alpha-helix frequency, calculated from the AAindex descriptor for normalized alpha-helix frequency and averaged across valid non-padding residues in each 31-residue peptide window. **(C)** Density distribution of isoelectric point, calculated from the AAindex isoelectric-point descriptor and averaged across valid non-padding residues in each 31-residue peptide window. **(D)** Density distribution of beta-turn frequency, calculated from the AAindex descriptor for normalized beta-turn frequency and averaged across valid non-padding residues in each 31-residue peptide window. **(E)** Density distribution of hydropathy index, calculated from the AAindex Kyte-Doolittle hydropathy descriptor and averaged across valid non-padding residues in each 31-residue peptide window. **(F)** Density distribution of B-value-derived flexibility, calculated from the AAindex descriptor for normalized flexibility parameters based on B-values and averaged across valid non-padding residues in each 31-residue peptide window. **(G)**Density distribution of hydrophilicity value, calculated from the AAindex Hopp-Woods hydrophilicity descriptor and averaged across valid non-padding residues in each 31-residue peptide window. For panels B-G, group differences were assessed using the two-sided Mann-Whitney U test with FDR correction, and the U statistic was converted to a signed Z-score to indicate the direction and magnitude of group differences.

Collectively, these results demonstrate that Deep-Palm not only accurately predicts S-palmitoylation sites, but also captures sequence, structural and physicochemical patterns associated with S-palmitoylation, suggesting that the biologically informed modeling strategies of Deep-Palm effectively uncover determinants for S-palmitoylation site recognition.

### Deep-Palm facilitates the discovery of novel S-palmitoylation sites from MS data

Current model training is based on database-annotated sites, limiting novel site discovery. To test tool’s ability to discover such sites, we assessed Deep-Palm on two independent biotin-labeled MS datasets for palmitoylation: a mouse hepatocellular carcinoma dataset^[38]^ and a human HeLa dataset^[39]^. We annotated S-palmitoylation sites based on the MS data as the positive cases. Non-palmitoylated sites were defined as sites detected in the matched proteome dataset but not identified as palmitoylated, thereby minimizing false negatives caused by insufficient protein expression. Using these datasets, we evaluated the performance of Deep-Palm. The model demonstrated robust discriminatory capability across both datasets, achieving AUC values of 0.822 in the mouse hepatocellular carcinoma dataset and 0.759 in the HeLa cell line dataset (Fig. 6A). Compared with existing predictors, Deep-Palm achieved the highest sensitivity toward MS-positive sites in both datasets, correctly recalling 89% of experimentally detected positive sites in the mouse hepatocellular carcinoma and HeLa datasets (Fig. 6B, C). The enhanced sensitivity of Deep-Palm toward MS-supported sites suggests its ability to discover potential true-positive S-palmitoylation events.

**Figure 6.**
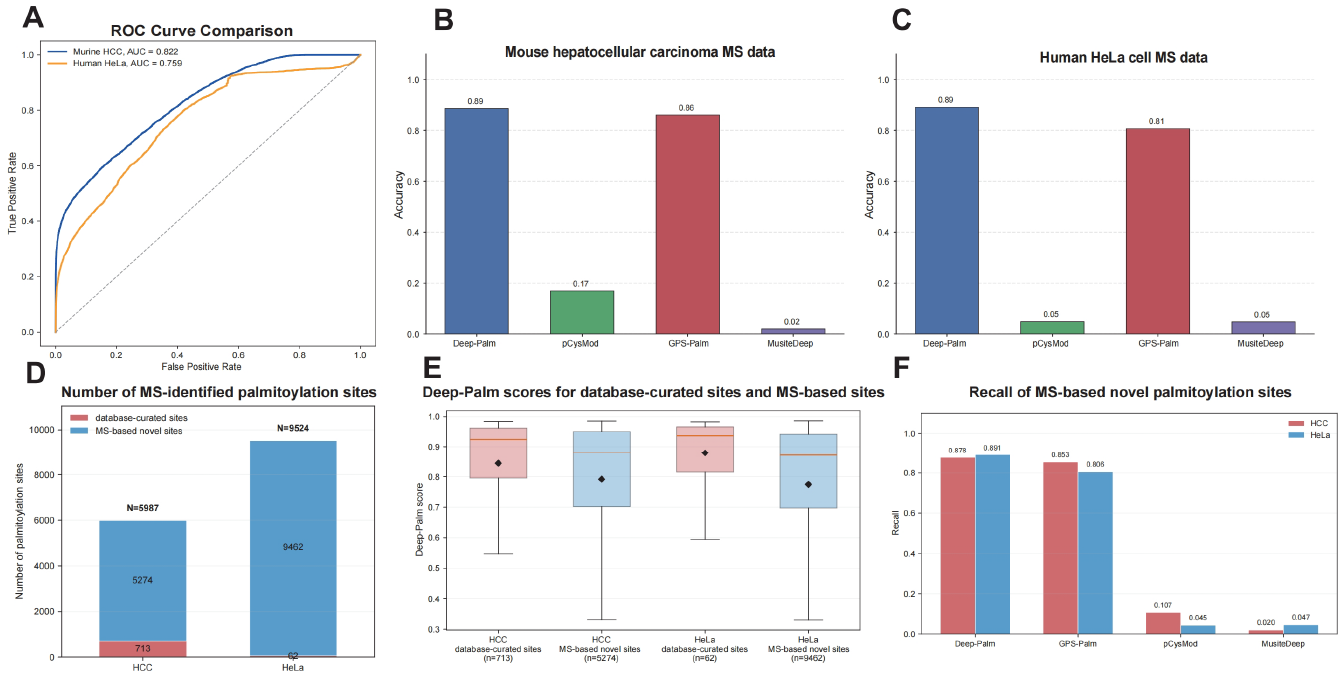
Independent MS-based evaluation of Deep-Palm for S-palmitoylation-site prediction. **(A)** ROC curve comparison of Deep-Palm predictions in the mouse hepatocellular carcinoma and human HeLa cell datasets using the subset with both positive and negative labels. **(B)** Positive-site accuracy of Deep-Palm, pCysMod, GPS-Palm, and MusiteDeep in the mouse hepatocellular carcinoma MS dataset. **(C)** Positive-site accuracy of the same methods in the human HeLa cell MS dataset. **(D)** Numbers of database-curated sites and MS-based novel sites among the MS-identified positive S-palmitoylation sites in the mouse hepatocellular carcinoma and human HeLa cell datasets. **(E)** Deep-Palm score distributions for database-curated sites and MS-based novel sites in the mouse hepatocellular carcinoma and human HeLa cell datasets. Boxes show the interquartile range, horizontal lines indicate medians, and black diamonds indicate means. **(F)** Recall of Deep-Palm, pCysMod, GPS-Palm, and MusiteDeep for MS-based novel sites in the mouse hepatocellular carcinoma and human HeLa cell datasets. In panels B, C, and F, only positive sites were evaluated; therefore, positive-site accuracy and recall represent the fraction of positive sites correctly predicted by each method.

To evaluate the ability of Deep-Palm to identify novel S-palmitoylation sites, we removed the sites that had already been annotated in public databases (all the training and testing sites used by the model) from the MS-positive set (Fig. 6D). Using the remaining palmitoylation sites identified by MS, Deep-Palm achieved only slightly reduced sensitivity compared with database-annotated sites, with sensitivities of 0.878 versus 0.931 in the mouse dataset and 0.891 versus 1.000 in the human dataset, respectively (Fig. 6E). Notably, among the compared methods, Deep-Palm still retained the highest sensitivity in both the mouse hepatocellular carcinoma and HeLa datasets (Fig. 6F). Collectively, these results suggest that Deep-Palm generalizes effectively to previously unreported experimentally detected sites and offers a cost-effective strategy to facilitate novel palmitoylation event discovery prior to MS validation.

## Discussion

Current experimental annotation of S-palmitoylation remains low-throughput and costly, highlighting the need for computational prediction tools. Several tools have been developed for S-palmitoylation-site discovery with cysteine-centered sequence^[12-17,40,41]^. These tools incorporate sequence-derived features such as motif, k-mer or BLAST-based similarity to known sites. To accurately model S-palmitoylation determination process in cell, it is essential to incorporate features that capture underlying biological and physicochemical properties. Thus, in this study, we developed Deep-Palm, a biologically-informed deep learning framework that integrates four multi-view branches: amino acid sequence, protein properties, ESM embedding and spatial structure.

On the independent test set, Deep-Palm achieved an AUC of 0.950 and showed accurate performance with individual branch. Among the four branches, protein properties branch achieved the highest AUC (0.939), indicating the important role of surrounding physicochemical context of cysteine in palmitoylation determination, and further supports the reliability of Deep-Palm’s biologically informed feature construction strategy. Feature analyses revealed distinct spatial structural and physicochemical differences between palmitoylated and non-palmitoylated peptide windows. Palmitoylated sites exhibited larger local residue-residue distances, suggesting a more open local structural environment. In addition, multiple physicochemical descriptors showed differences between the two groups, including α-helix frequency, hydrophobicity and pI. These results indicate that S-palmitoylation is associated with coordinated structural and physicochemical contexts, which are effectively captured by Deep-Palm.

Deep-Palm captures key biological determinants underlying S-palmitoylation-site recognition, enabling consistently accurate performance across diverse protein types and sequence contexts. When the testing set was stratified according to cysteine density, nearest-neighbor distance and the relative N- to C-terminal position of the target cysteine, Deep-Palm maintained stable performance across all stratifications and was better than existing predictors. Further evaluation across Gene Ontology categories showed consistently high AUC values (>0.90). Together, these results demonstrate that Deep-Palm is robust to both local sequence heterogeneity and functional annotation diversity.

As a computational method for S-palmitoylation identification, a key limitation of Deep-Palm is that the model relies on current annotations, which may be biased by database curation methods, experimental technology and protein expression abundance. However, validated on two independent MS datasets, Deep-Palm achieved robust performance (AUC 0.822 and 0.759) and recalled 89% of MS-positive sites which are absent from public databases. Thus, Deep-Palm provides a cost-free, computation-only strategy to discover novel S-palmitoylation sites prior to experimental validation. Collectively, Deep-Palm integrated multi-view features for S-palmitoylation prediction. It not only accurately identifies *bona fide* S-palmitoylation cysteines but also provides insights into the palmitoylation site recognition pattern within cells, facilitating the understanding of palmitoylation mechanisms in multiple biological processes and diseases.

## Acknowledgements

This work was supported by the National Natural Science Foundation of China (Grant No. 32500579), the Chongqing Key program for special projects of technological innovation and application development (CSTB2023TIAD-KPX0050), and the Fundamental Research Funds for the Central Universities (Project No.2025CDJZKPT-10).

## Author contributions

B.X., Y.K. and M.D. designed the project. M.D., S.F. and W.W. developed the prediction model. J.H. and H.W. analyzed the mass spectrometry data. M.D. and L.L. contributed to the model interpretation analyses. Y.K. provided advice on statistical analysis. M.D., Y.K. and B.X. wrote the manuscript. B.X. and Y.K. supervised the study.

## Data and code availability

The processed datasets, source code, pretrained Deep-Palm model and input feature files required to reproduce the model training and prediction workflows are available at https://github.com/DML666666/Deep-Palm. The Deep-Palm predicted S-palmitoylation sites for human and mouse proteins can be accessed via https://palmlab.intelligent-oncology.com/.

## Competing interests

All the authors declare no competing interests.

## Notes

### Competing Interest Statement

The authors have declared no competing interest.

### Summary of Updates

This revised manuscript provides an updated and more rigorous evaluation of Deep Palm. The independent test analysis was updated and the reported AUC of the final model changed from 0.931 in the previous version to 0.950 in the revised version. The introduction was revised to better position the study in relation to existing S palmitoylation predictors and protein language model based methods. The methods were expanded to clarify dataset construction feature calculation ensemble learning statistical analysis mass spectrometry based site identification and comparison with existing predictors. The results and figure descriptions were updated to report performance across cysteine sequence contexts and Gene Ontology functional groups and to strengthen interpretation of spatial structure and protein property features. Evaluation using the mouse hepatocellular carcinoma dataset and the human HeLa cell dataset was clarified to assess the identification of previously unannotated candidate sites. Author affiliations author contribution information references terminology and data availability information were also updated.

